# Macrovesicular steatosis in nonalcoholic fatty liver disease is a consequence of purine nucleotide cycle driven fumarate accumulation

**DOI:** 10.1101/2020.06.23.166728

**Authors:** Matthew C. Sinton, Baltasar Lucendo Villarin, Jose Meseguer Ripolles, Sara Wernig-Zorc, John P. Thomson, Paul D. Walker, Alpesh Thakker, Gareth G. Lavery, Christian Ludwig, Daniel A. Tennant, David C. Hay, Amanda J. Drake

**Affiliations:** University/BHF Centre for Cardiovascular Science, The Queen’s Medical Research Institute, Edinburgh BioQuarter, 47 Little France Crescent, Edinburgh EH16 4TJ; Centre for Regenerative Medicine, Institute for Regeneration and Repair, Edinburgh BioQuarter, 5 Little France Crescent, Edinburgh, EH16 4UU; Department of Biochemistry, University of Regensburg, Universitätsstraße 31, 93053 Regensburg, Germany; Human Genetics Unit, MRC Institute for Genetics and Molecular Medicine, University of Edinburgh, Crewe Road South, Edinburgh, EH4, 2XU; Institute of Metabolism and Systems Research, IBR Tower, College of Medical and Dental Sciences, Edgbaston, University of Birmingham, Birmingham, B15 2TT

**Keywords:** Nonalcoholic fatty liver disease, steatosis, TCA cycle, mitochondria, purine nucleotide cycle, fumarate, succinate

## Abstract

Nonalcoholic fatty liver disease (NAFLD) affects ~88% of obese individuals and is characterised by hepatic lipid accumulation. Mitochondrial metabolic dysfunction is a feature of NAFLD. We used a human pluripotent stem cell-based system to determine how mitochondrial dysfunction is linked to hepatic lipid accumulation. We induced lipid accumulation in hepatocyte-like cells (HLCs) using lactate, pyruvate and octanoate (LPO). Transcriptomic analysis revealed perturbation of mitochondrial respiratory pathways in LPO exposed cells. Using ^13^C isotopic tracing, we identified truncation of the TCA cycle in steatotic HLCs. We show that increased purine nucleotide cycle (PNC) activity fuels fumarate accumulation and drives lipid accumulation in steatotic cells. These findings provide new insights into the pathogenesis of hepatic steatosis and may lead to an improved understanding of the metabolic and transcriptional rewiring associated with NAFLD.

## Introduction

Nonalcoholic fatty liver disease (NAFLD) is the most common form of hepatic disease and is strongly associated with obesity and the development of type 2 diabetes (WHO, 2006). In the earliest stage of NAFLD, triglyceride (TG) accumulation in hepatocytes leads to the development of hepatic steatosis (Valenti et al., 2016). This is characterised by the development of macrovesicular steatosis, whereby TGs are stored in large lipid droplets (Wang and Yu, 2016). Whilst steatosis is largely benign, it can progress to nonalcoholic steatohepatitis (NASH), which, in turn, increases the risk of developing cirrhosis and hepatocellular carcinoma (Asrih and Jornayvaz, 2015). However, the mechanism(s) underlying this progression have yet to be determined. At present, there are no specific therapeutics available to reverse or treat NAFLD, and the only effective intervention is through the reduction of obesity following bariatric surgery (Laursen et al., 2019).

NAFLD has been linked to impaired mitochondrial respiration, with previous findings revealing altered electron transport chain (ETC) activity associated with steatosis (Koliaki et al., 2015; Sinton et al., 2020). Rodent studies suggest that steatosis is preceded by mitochondrial dysfunction, indicating that NAFLD and its progression may, in part, be a consequence of impaired mitochondrial respiration (Rector et al., 2010). Further, hepatic steatosis in mice is accompanied by increased tricarboxylic acid (TCA) cycle activity (Satapati et al., 2012). Human studies also suggest that NAFLD is associated with increased TCA cycle activity, with increased movement of substrates into and out of the TCA cycle alongside increased rates of gluconeogenesis and lipolysis (Sunny et al., 2011). However, whilst informative, these studies were indirect and used measurements of plasma metabolites to infer hepatic metabolic flux, rather than directly measuring levels within the liver. Examination of TCA cycle function in mouse models of NAFLD identified increased hepatic TCA cycle flux (Satapati et al., 2015). Limiting pyruvate transport into the mitochondria reduces mitochondrial flux and limits the development of NASH-associated inflammation (Rauckhorst et al., 2017). However, metabolic flux analysis in whole tissue is confounded by the presence of heterogeneous cell populations, each of which may have a different metabolic phenotype (Antoniewicz, 2018). Studies in primary rat hepatocytes show that modulation of TCA cycle anaplerosis fuels oxidative stress and enhances development of a NASH phenotype (Egnatchik et al., 2019). One limitation of primary cells is that they rapidly lose their phenotype and display altered transcriptional signatures on purification (Godoy et al., 2016; Knobeloch et al., 2012). In these studies, we employed a renewable form of human liver tissue (Wang et al., 2019b). To our knowledge, high-resolution metabolomic analyses have not been performed in a renewable human system, providing the field with new insight into human liver disease.

The TCA cycle metabolites alpha-ketoglutarate (αKG), succinate and fumarate are allosteric regulators of the αKG-dependent dioxygenase family of enzymes; their activity is promoted by αKG and inhibited by succinate and fumarate (Hausinger, 2004; Roach et al., 1995; Xiao et al., 2012). The Fe(II)/O_2_-dependent enzymes include the Jumonji domain-containing histone demethylases (JHDMs), prolyl-4-hydroxylases, and the ten-eleven translocation (TET) dioxygenases (Laukka et al., 2016; McDonough et al., 2010). The JHDMs and TET enzymes can regulate the epigenome through modulation of histone and DNA methylation, respectively, with the latter oxidising 5-methylcytosine to 5-hydroxymethylcytosine (5hmC) (Pollard et al., 2008; Tahiliani et al., 2009). In human cancers, mutations in succinate dehydrogenase and fumarate hydratase result in the accumulation of succinate and fumarate, which inhibit JHDM and TET dioxygenase activity and may contribute to tumourigenesis (Xiao et al., 2012). Dysregulation of the αKG-dependent dioxygenase enzymes may also be important in the pathogenesis of liver disease; we have previously identified alterations in 5hmC at specific loci in murine and human models of NAFLD, suggesting a role for the TET enzymes, which oxidise 5-methylcytosine (5mC) to 5-hydroxymethylcytosine (5hmC) (Lyall et al., 2020). Studies in mouse models of hepatocellular carcinoma suggest that the TET enzymes may also contribute to the progression of liver cancer, through perturbation of normal DNA demethylation events at promoters (Thomson et al., 2016).

In this study, we aimed to understand the impact of lipid accumulation on TCA cycle activity and whether this influences TET enzyme activity. We performed these studies in a reliable model of NAFLD (Lyall et al., 2018) which enables us to repeatedly study hepatocyte biology in isolation, avoiding the confounding effects of tissue disaggregation, bulk tissue analysis or inferred measurement.

## Results

### LPO treatment induces intracellular lipid accumulation and transcriptomic alterations in key mitochondrial respiratory pathways

HLC exposure to LPO resulted in macrovesicular steatosis (Fig. 1A), as previously reported (Sinton et al., 2020). This was associated with widespread transcriptomic changes (Fig. 1B). Differential gene expression analysis identified 853 downregulated and 826 upregulated genes (log_2_ fold change cut-off >1.5) in LPO exposed cells compared with control. A selection of candidate genes was validated by RT-qPCR (Fig. S2). From the RNA-seq analysis, we observed altered expression of a number of genes previously described as having function roles in the progression of NAFLD, including PLIN2, PPARGC1A, CYP7A1, and HMGCS2 (Fig. 1C). Mapping genes with a log_2_ fold change >1.5 to the KEGG pathway database identified a number of enriched pathways (Fig. 1D & 1E), including those related to steroid hormone biosynthesis and ascorbate and aldarate metabolism, containing 18 and 7 genes respectively (Table S1). There was extensive downregulation of the histone structural units H1, H2A/B, H3 and H4 and UDP-glucuronosyltransferases, which were enriched in multiple pathways. Specifically comparing transcriptomic data with the KEGG terms ‘TCA Cycle’ and ‘Oxidative Phosphorylation’ revealed extensive gene expression changes within these pathways (Fig. 1F & 1G) including in the majority of genes encoding enzymes that catalyse metabolite interconversion (Table S2). Analysis of the Oxidative Phosphorylation pathway revealed an overall downregulation of transcription of genes encoding components of respiratory complexes I (ND1, ND2, ND4L, NDUFS2, NDUFV1, NDUFA10, NDUFB2 NDUFA2, NDUFB10) and IV (COX1, COX4I1, COX8A, COX6B1), as well as ATP synthase (ATP6V1A, ATP5MF, ATP6V0D1, ATP6V1E1, ATP6, APT5MC2, ATP5MC1) (Table S3).

**Figure 1.**
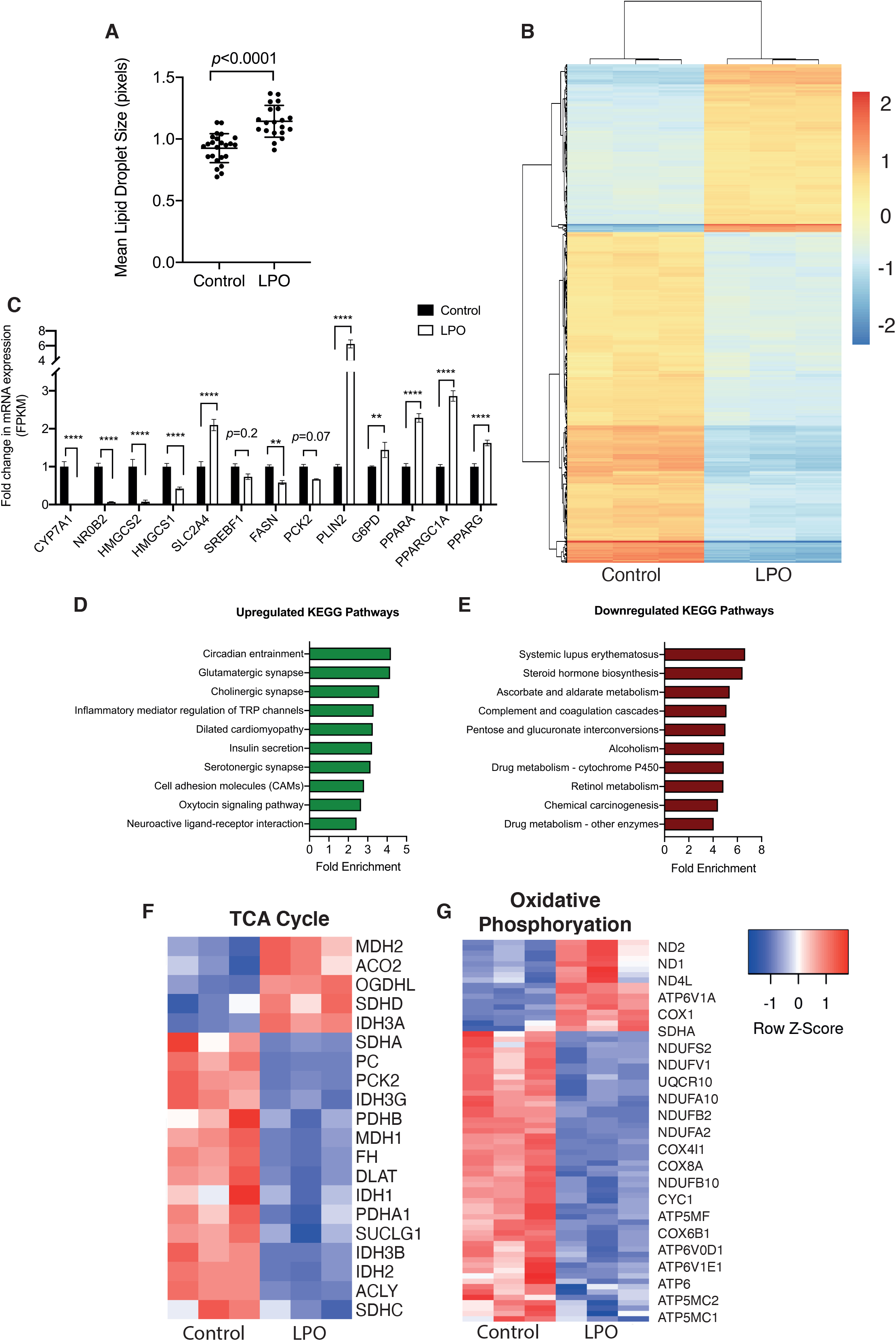
LPO treatment induces macrovesicular steatosis in HLCs, which is associated with transcriptional rewiring reflecting human NAFLD. (**A**) Intracellular lipid droplets increase in size in response to LPO (*n* = 24 (control) and 20 (LPO) biological replicates/group). (**B**) Heatmap analysis of transcriptional changes associated with macrovesicular steatosis. (**C**) Expression of key NAFLD-associated genes is disrupted in steatotic HLCs. (**D-E**) Pathway enrichment analysis reveals a number of disrupted pathways associated with macrovesicular steatosis. (**F-G**) Analysis of KEGG TCA cycle and oxidative phosphorylation pathways reveals extensive disruption of expression of key genes. Data in (**A**) were analysed using a two-tailed Student’s t-test. Data in (**C**) were analysed by two-way ANOVA with Sidak post-hoc testing. Unless otherwise specified, *n* = 3 biological replicates per group. Data are expressed as mean ± SD.

### Induction of a NAFLD-like phenotype in HLCs is associated with increased pyruvate carboxylase activity

To investigate TCA cycle dynamics, lactate in the LPO cocktail was replaced with ^13^C_3_-lactate. Lactate can enter the TCA cycle *via* mitochondrial pyruvate metabolism, through either pyruvate carboxylase (PC) or pyruvate dehydrogenase (PDH). Alternatively, pyruvate can be transported into the cytosol and used in gluconeogenesis (Fig. 2A). LPO exposure was associated with increases in steady state levels of pyruvate, aspartate and citrate, but there were no changes in metabolites associated with the gluconeogenesis pathway (Fig. 2B). Although pyruvate was added as part of the LPO cocktail, increased pyruvate generation also occurred as a result of increased lactate dehydrogenase activity (Fig. 2C). Although we were unable to directly measure oxaloacetate levels, we used aspartate as a surrogate and found increased m+3 labelling, demonstrating direct conversion between these two metabolites (Fig. 2D). Furthermore, isotopomer labelling of citrate indicated preferential synthesis from oxaloacetate and decreased conversion of labelled acetyl-CoA by PDH (Fig. 2E). An alternative pathway for pyruvate is conversion to alanine and whilst there were no changes in steady state alanine levels, there were moderate, but significant, changes in the incorporation of ^13^C (Fig. 2F), indicating sustained interconversion of these two metabolites. Despite there being no increase in steady state levels of gluconeogenesis-associated metabolites, the increased m+2 labelling of 3-PG (Fig. 2G), serine (Fig. 2H), and glycine (Fig. 2I), suggests increased flux through the gluconeogenesis pathway.

**Figure 2.**
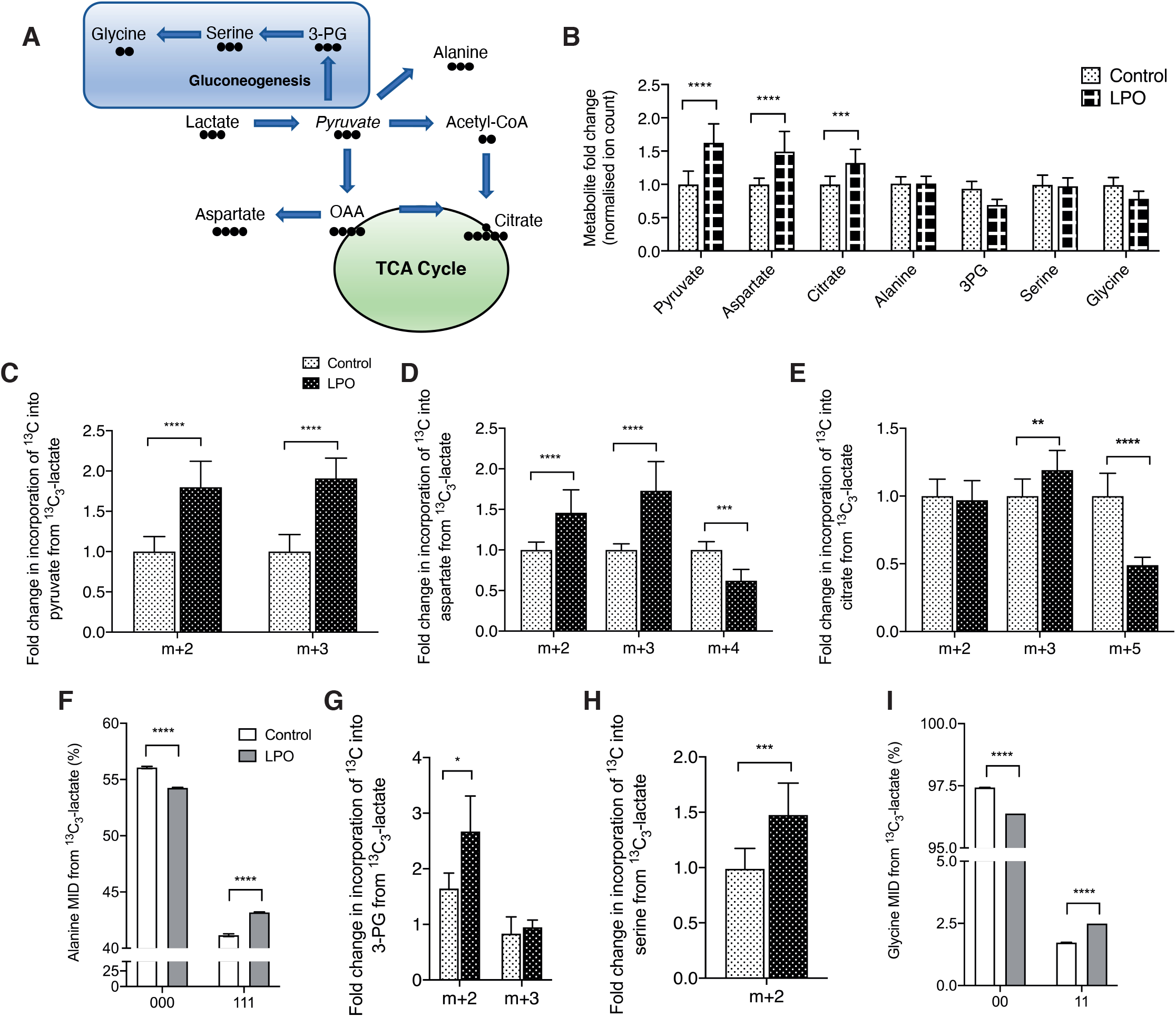
Macrovesicular steatosis is associated with increased PC activity, leading to preferential anaplerosis of pyruvate into the TCA cycle. (**A**) Schematic outlining conversion of lactate to pyruvate and routes by which this can be converted. Black circles denote carbon atoms. (**B**) Steady state measurements of pyruvate and metabolites that it can be converted to. Isotopomer labelling patterns of pyruvate (**C**), aspartate (**D**) and citrate (**E**) show anaplerosis into the TCA cycle. There is minimal conversion of pyruvate to alanine (**F-G**), but increased flux of the m+2 isotopomer through metabolites related to gluconeogenesis, with sustained conversion to 3-PG, serine and glycine (**H-K**). For NMR data, labels are: 1 = ^13^C labelling; 0 = no ^13^C labelling. All GC-MS data consisted of 10 biological replicates and 2 technical replicates. Isotopomer data were calculated by multiplying MID (multiple ion detection) by normalised total ion count. For NMR data (**G** and **K**) *n* = 4 biological replicates/group. Data were analysed by two-way ANOVA with Sidak post-hoc testing, or (**I-J**) two-tailed Student’s t-test. Data are expressed as mean ± SD.

### Steatosis in HLCs is associated with truncation of the TCA cycle

Isotopic labelling of TCA metabolites can produce a number of different isotopomers, dependent on the directionality of metabolite synthesis (Fig. 3A). In steatotic HLCs, we observed increases in steady state levels of the TCA cycle metabolites αKG, fumarate and malate (Fig. 3B). In contrast, there were no changes in the steady state levels of succinate and no evidence for increased cataplerosis through glutamate. To further analyse alterations in TCA cycle dynamics we measured ^13^C incorporation into metabolites. Whilst there was increased ^13^C incorporation into PC-derived citrate (Fig. 2E), this was not the case for αKG (Fig. 3C). The decreased levels of ^13^C incorporation into glutamate demonstrates that this does not result from increased cataplerosis through glutamate (Fig. 3D). The increased expression of OGDHL and decreased expression of SUCG1 (Table S2) suggest that steatosis is associated with the generation of an increased pool of succinyl-CoA, with impaired conversion to succinate. This is supported by substantially decreased incorporation of ^13^C into succinate (Fig 3E). In contrast, we observed increased ^13^C incorporation into fumarate, malate, and aspartate (Fig 3F & 3G). Since malate dehydrogenase and fumarate hydratase readily reverse their reactions (Dasika et al., 2015; Tyrakis et al., 2017), increased incorporation of ^13^C into both malate and fumarate indicate a reversal of TCA cycle activity. Taken together, this shows that truncation of the TCA cycle occurs in steatotic cells, with inhibition of the conversion of succinate to fumarate (Fig. 3H). The increased incorporation of ^13^C into oxaloacetate (using aspartate as a surrogate), demonstrates increased PC activity, driving the conversion of pyruvate to oxaloacetate and suggests possible disruption of associated metabolic cycles, including the malate-aspartate shuttle (MAS) and the purine nucleotide cycle (PNC).

**Figure 3.**
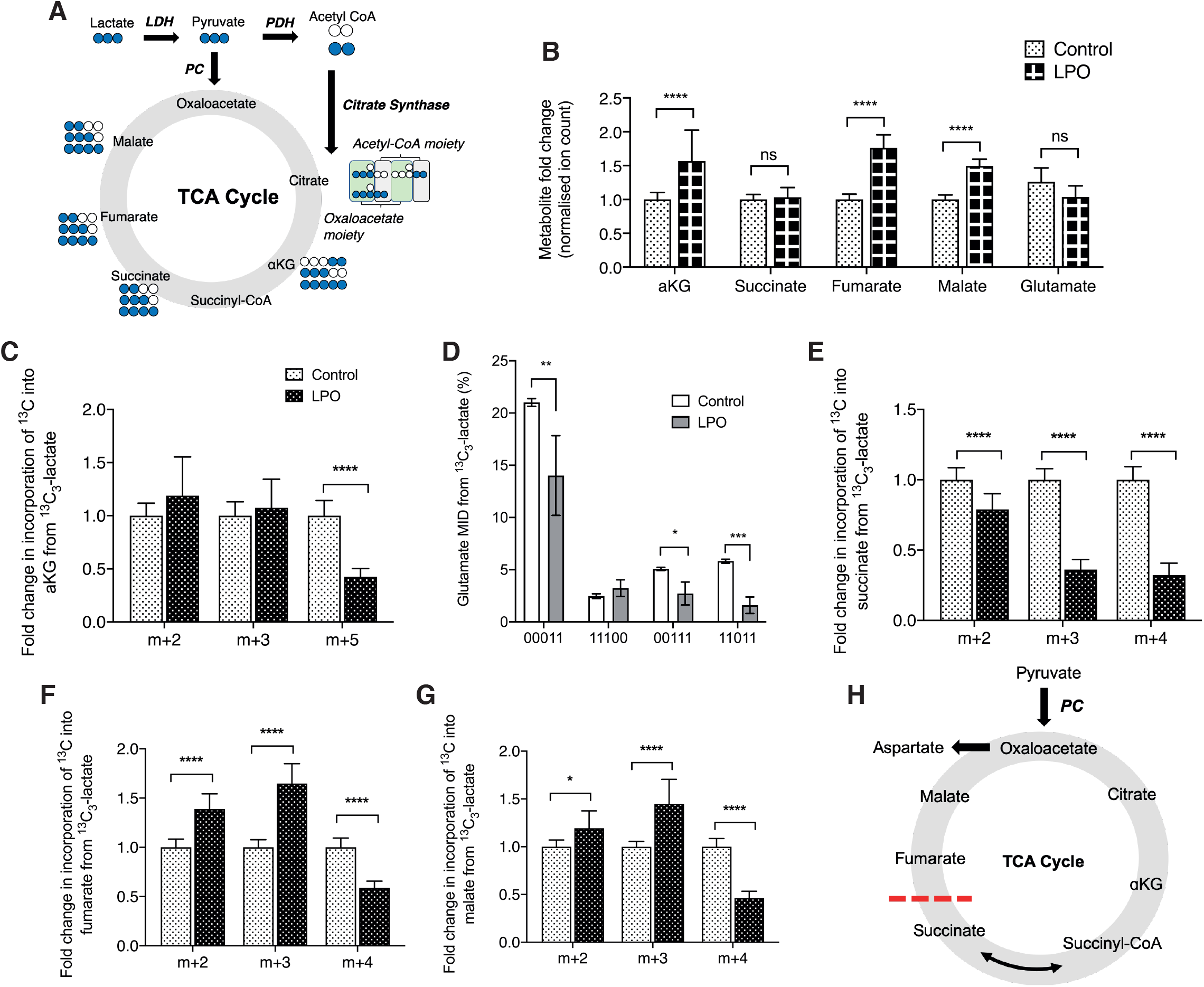
Macrovesicular steatosis in HLCs results in truncation of the TCA cycle, inhibiting conversion of succinate to fumarate. Paradoxically, this is associated with increased accumulation of fumarate. (**A**) Schematic outlining the TCA cycle and possible ^13^C labelled isotopomers from a single cycle. (**B**) Steady state measurements of TCA cycle-associated metabolites that could be measured by GC-MS. (**C**) PC-derived αKG is unchanged in the presence of steatosis, and PDH-derived αKG is reduced. (**D-E**) Labelling patterns of αKG are not due to increased cataplerosis through glutamate, as both PC- and PDH-derived glutamate are reduced in response to steatosis. (**F**) Furthermore, reduced ^13^C labelling of succinate indicates inefficient conversion from αKG. (**G-H**) Despite reduced labelling of succinate, both fumarate and malate show increased PC-derived label incorporation. (**I**) Proposed model of preferential anaplerosis of pyruvate into the TCA cycle, with concomitant truncation, preventing conversion of succinate to fumarate. For NMR data, labels are: 1 = ^13^C labelling; 0 = no ^13^C labelling. All GC-MS data consisted of 10 biological replicates and 2 technical replicates. Isotopomer data were calculated by multiplying MID (multiple ion detection) by normalised total ion count. For NMR data (**E**) *n* = 4 biological replicates/group. Data were analysed by two-way ANOVA with Sidak post-hoc testing and are expressed as mean ± SD.

### Increased PNC and MAS activity drives NAFLD-associated fumarate accumulation

Next, we wanted to determine the source of fumarate accumulation in steatotic HLCs. Transcriptomic analysis identified dysregulation of multiple genes associated with the MAS (Fig. 4A) and PNC (Fig. 4B), indicating that these pathways may be involved in fumarate accumulation. To investigate this further, HLCs incubated with ^13^C_3_-lactate-labelled LPO were co-incubated with either 5-Aminoimidazole-4-carboxamide-1-β-D-ribofuranosyl 5′-monophosphate (AICAR) or O-(Carboxymethyl)hydroxylamine hemihydrochloride (AOA), to inhibit the PNC or MAS, respectively. Addition of AICAR increased only the steady state levels of αKG, and neither AICAR nor AOA had a significant impact on the synthesis of pyruvate, citrate, succinate or malate (Fig. 4C). However, both inhibitors significantly reduced steady state levels of fumarate and aspartate. Whilst addition of AICAR did not impact on the synthesis of PDH-derived citrate (Fig. 4D), it did result in increased accumulation of PDH-derived ^13^C in αKG (Fig. 4E), suggesting impaired conversion to succinyl-CoA or increased anaplerosis from glutamate. AICAR was also able to partially restore the effects of steatosis on succinate levels (Fig. 4F), resulting in small but significant increases in PC- and PDH-derived isotopomers. The most profound effect of AICAR was on fumarate, with each isotopomer reduced to levels below those observed in the control group (Fig. 4G), demonstrating that in steatotic HLCs, fumarate accumulation is primarily driven by the PNC. Following AOA treatment, we observed a moderate decrease in PC-derived ^13^C incorporation into fumarate, demonstrating a small contribution from the MAS. AICAR had a moderate impact on incorporation of PC-derived incorporation of ^13^C into malate, whereas AOA impacted on both PC- and PDH-derived ^13^C incorporation, suggesting limited contribution of the MAS and PNC to malate accumulation (Fig. 4H). Both AICAR and AOA reduced incorporation of ^13^C into aspartate (Fig. 4I). These data show that in the presence of TCA cycle truncation, mitochondria increase both MAS and PNC activity (Fig. 4J).

**Figure 4.**
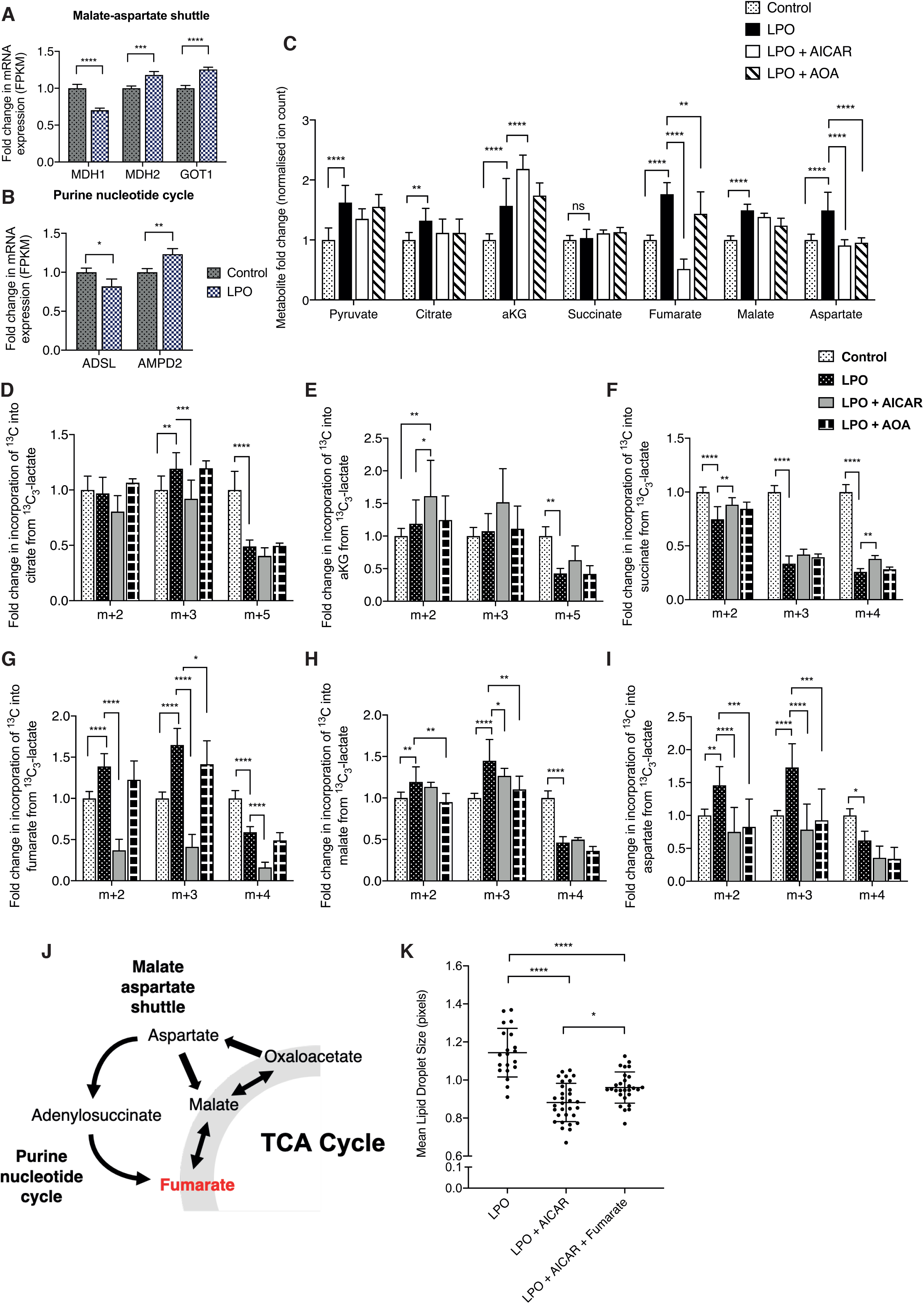
In the presence of TCA cycle truncation, the purine nucleotide cycle and malate-aspartate shuttle fuel fumarate accumulation. (**A-B**) Transcriptomic analysis showed increased expression of malate-aspartate shuttle (MAS) and purine nucleotide cycle (PNC) transcripts, indicating perturbed activity. (**C**) Inhibition of the PNC and MAS reversed steatosis-induced accumulation of fumarate and aspartate, but not malate. (**D**) PNC inhibition moderately impacted PC-derived citrate ^13^C labelling, but not (**E**) αKG or (**F**) succinate. (**G**) In contrast, PNC inhibition profoundly reduced incorporation of ^13^C into fumarate. MAS inhibition also limited PC-derived fumarate accumulation, but to a lesser extent. (**H**) PNC and MAS expression also inhibited generation of PC-derived malate, but to a much smaller extent than fumarate. (**I**) PNC and MAS inhibition also resulted in a significant reduction in the generation of PC-derived aspartate. (**J**) Schematic outlining the proposed pathways by which fumarate accumulation occurs in response to steatosis. (**K**) Inhibition of fumarate through the PNC reduced LPO-induced macrovesicular steatosis, which was partially restored through addition of exogenous monomethyl fumarate. For NMR data, labels are: 1 = ^13^C labelling; 0 = no ^13^C labelling. AICAR = PNC inhibitor; AOA = MAS inhibitor. All Control and LPO group GC-MS data consisted of 10 biological replicates and 2 technical replicates, as shown in Figure 3. GC-MS LPO + AICAR and LPO + AOA groups consisted of 6 biological replicates/group. Isotopomer data were calculated by multiplying MID by normalised total ion count. For lipid droplet analysis (**K**), the LPO group is as shown in Figure 1. For the LPO + AICAR and the LPO + AICAR + Fumarate group, *n* = 31 and 28 biological replicates/group, respectively. Data were analysed by two-way ANOVA with Sidak post-hoc testing and are expressed as mean ± SD.

### Inhibition of fumarate accumulation inhibits lipid droplet hypertrophy in LPO-treated HLCs

To determine whether manipulation of fumarate levels affected the development of macrovesicular steatosis, we performed HCA microscopy on HLCs. Cells treated with LPO and AICAR did not develop macrovesicular steatosis (Fig. 4K). The addition of exogenous fumarate to steatotic HLCs treated with AICAR resulted in the development of larger lipid droplets.

### Fumarate accumulation is not associated with widescale alterations of 5hmC in protein-coding regions in steatotic HLCs

As we observed increased synthesis of fumarate in steatotic HLCs, we aimed to determine whether this would impact on TET enzyme activity, as determined by changes in 5hmC. To do so we carried out genome-wide 5hmC sequencing, identifying 3294 differentially hydroxymethylated regions (DHRs) between LPO exposed and control cells (>2-fold change, see methods). Within each group, samples were highly correlated, suggesting that the 5hmC patterns are stable between replicate samples (Fig. 5A), with the majority of change located in intragenic or intronic regions (Fig. 5B). It is thought that 5hmC in gene body regions may functionally relate to mRNA transcription (Thomson et al., 2016). Therefore, we generated heatmaps representing mean changes across the gene body, with cluster 1 showing subtle increases in 5hmC within the TSS region (Fig. 5C). Analysis of genic 5hmC patterns reveal that a number of promoter regions display changes in 5hmC levels following LPO exposure. (Fig. 5D). In line with this, we integrated DHR and RNA-seq data to look at promoter regions upstream of the transcriptional start site. In doing so, we identified 12 promoter regions with differential enrichment of 5hmC (>2-fold) and where mRNA expression was altered >0.5-fold. Linear regression of these regions identified a moderate but significant negative relationship between 5hmC enrichment and mRNA expression (Fig. S1; Table S4). Of these genes 7 (EPHX3, ERO1B, CSGALNACT1, DOC2A, COL6A1, CASP1 and TMEM88) were previously shown to be dysregulated in the pathogenesis of NAFLD or progression to cirrhosis (Atanasovska et al., 2017; Cazanave et al., 2017; Parafati et al., 2018; Revill et al., 2013; Wilson and Kumar, 2018). This may indicate a role for 5hmC in the regulation of these genes in NAFLD pathogenesis.

**Figure 5.**
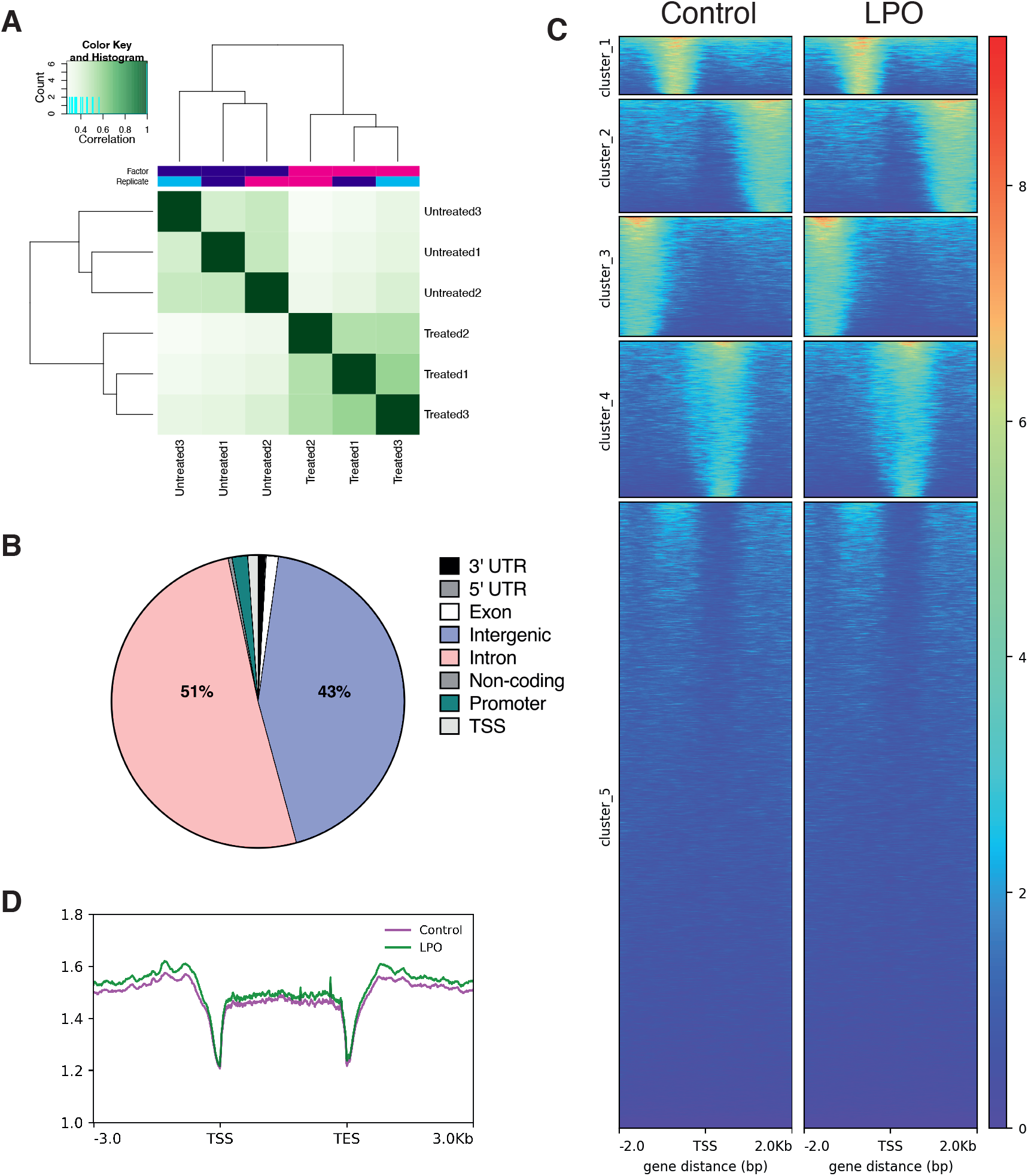
Macrovesicular steatosis in HLCs does not correlate with wide-scale changes in 5hmC enrichment. (**A**) Correlation heatmap of control vs. LPO groups, following hmeDIP sequencing, clustered by Euclidean distance. (**B**) Proportion of DHRs associated with different regions of the genome, showing the majority located in intragenic and intronic regions. (**C**) Heatmap of DHRs, with an FDR of <0.05 and k-means clustering. (**D**) Sliding window analysis of the transcriptional start site (TSS), gene body, and transcriptional end site (TES) shows minimal changes between control and LPO groups. For both control and LPO, *n* = 3 biological replicates/group.

## Discussion

NAFLD is a challenging disease to study in humans. This is due to the difficulty of obtaining tissues that are not compromised by confounding conditions which make high resolution analysis of transcriptional and metabolic rewiring difficult. Using this HLC-based model of NAFLD we were able to overcome many of these difficulties.

Through transcriptomic analysis, we observed disruption of the expression of genes in multiple pathways related to metabolism, confirming the findings of a number of previous studies in humans and mouse (Collison et al., 2009; Hardwick et al., 2013; Kolwankar et al., 2007; Liu et al., 2016; Nikolaou et al., 2019; Schiöth et al., 2016; Suppli et al., 2019; Yamaguchi and Murata, 2013). Previous studies in mouse models and indirect studies in humans have reported increases in hepatic TCA cycle activity associated with NAFLD (Satapati et al., 2012, 2015; Sunny et al., 2011). Additionally, compromised ETC activity with decreased mitochondrial maximal respiration has been reported in response to hepatic steatosis (Koliaki et al., 2015; Sinton et al., 2020). However, it was unclear whether impaired maximal respiration is solely due to downregulation of respiratory chain units or whether this is, in part, due to chemical inhibition of complexes I-III by octanoic acid, as previously described in rat liver (Scaini et al., 2012). Our transcriptomic analysis revealed substantial disruption of TCA cycle and oxidative phosphorylation pathways, indicating compromised energy metabolism in HLCs. In particular, our findings that transcription of multiple subunits of respiratory complexes I, IV and ATPase were downregulated suggest that transcription-based disruption of the ETC plays a role in impaired respiration.

Traditionally, NAFLD is also thought to increase TCA cycle activity (Satapati et al., 2012; Sunny et al., 2011) but, to date, detailed information regarding flux dynamics has remained elusive. We utilised stable isotopic tracing, using ^13^C_3_-lactate which provides greater metabolite labelling than other substrates (Hui et al., 2017). In the liver, flux of substrates into the TCA cycle predominantly occurs *via* PC (Lardy et al., 1965), and there is evidence suggesting that PC activity increases in NAFLD (Sunny et al., 2011), and contributes to increased rates of gluconeogenesis through conversion of oxaloacetate to phosphoenolpyruvate (Satapati et al., 2012). The data presented here support the assertion that PC activity increases in steatotic cells. However, while we observed sustained synthesis of gluconeogenesis-associated metabolites occurring in steatotic HLCs, our data show no associated increase in gluconeogenesis. Increased incorporation of only two ^13^C atoms into each of these metabolites may suggest that pyruvate-derived acetyl-CoA is being converted back to pyruvate *via* ketogenesis before entering gluconeogenesis. Increased acetyl-CoA pools and disrupted ketogenesis have been observed in NAFLD (Fletcher et al., 2019), lending support to this assertion. This demonstrates that pyruvate is predominantly being utilised to sustain TCA cycle activity in steatotic HLCs.

The increased flux of substrates into the TCA cycle also leads to higher levels of TCA cycle activity, as previously reported (Sunny et al., 2011). However, our data show that changes in TCA cycle are more nuanced that was previously suggested. We identified a truncation of the TCA cycle, with inhibition of the conversion of succinate to fumarate. This raised the question of why less ^13^C is being incorporated into succinate. The lack of cataplerosis through glutamate suggests that this is due to diminished conversion of succinyl-CoA to succinate and generation of an increased pool of succinyl-CoA. This is supported by the observation of decreases in PC-derived glutamate and succinate, despite no changes in PC-derived αKG. Since interconversion between succinyl-CoA and succinate is in near equilibrium and readily reversible (Lynn and Guynn, 1978), we propose that this effect could be a result of diminished succinyl-CoA synthetase activity.

Despite truncation of the TCA cycle, we found that HLCs are able to rewire their metabolic circuitry to compensate for this, generating increased levels of fumarate, predominantly through the PNC, and to a lesser extent the MAS. A similar metabolic bypass has been reported in cardiac ischemia (Chouchani et al., 2014), as well as human and mouse *in vitro* models of tumorigenesis (Tyrakis et al., 2017). In ischemic reperfusion injury, increased PNC activity results in fumarate overflow, driving reversal of succinate dehydrogenase activity and accumulation of succinate (Chouchani et al., 2014). We did not observe the same phenomenon here, but rather found evidence for inhibition of succinate dehydrogenase activity in steatotic HLCs. Although it is possible for fumarate to be generated from reversal of fumarate hydratase (FH) activity (Chouchani et al., 2014), we were unable to directly manipulate the activity of this enzyme in order to assess its contribution to the fumarate pool in steatotic HLCs. However, as FH operates at equilibrium (Ajalla Aleixo et al., 2019) it is possible that PNC-fuelled fumarate accumulation prevents reverse catalytic activity. Defects in complex I can lead to a reduction in the levels of NAD^+^ (Porcelli et al., 2010) and regeneration of fumarate *via* the PNC may be one mechanism of increasing the NADH pool and maintaining the hydrogen ion gradient of the ETC. The synthesis of fumarate by the PNC also results in the generation of AMP, which in turn is deaminated to produce ammonia (Arinze, 2005). In rodents, hyperammonaemia is associated with the progression of hepatic steatosis to cirrhosis (De Chiara et al., 2020), and our findings suggest that this is generated through increased PNC activity. This leads to the question of whether NAFLD-associated cirrhosis is then a consequence of ammonia production or a downstream effect of fumarate accumulation.

A further consequence of fumarate accumulation is the development of macrovesicular steatosis. Inhibition of the PNC prevented development of macrovesicular lipid droplets, and this was partially restored following the addition of exogenous fumarate. This correlates with findings in oligodendrocytes and CD8+ T cells, in which exposure to exogenous fumarate resulted in perturbed lipid metabolism (Bhargava et al., 2019; Huang et al., 2015), although the mechanism by which this occurs remains unknown. However, our data suggest that fumarate is important for the development of macrovesicular steatosis in HLCs and that mechanisms compensating for TCA cycle truncation in LPO-treated HLCs cells drive intracellular lipid accumulation.

Another potential consequence of fumarate accumulation in steatotic HLCs was inhibition of αKG-dependent dioxygenase enzyme activity. Given our previous data showing altered 5hmC in a mouse model of NAFLD and the potential importance of the TET enzymes in hepatocellular carcinoma (Lyall et al., 2020; Thomson et al., 2016), we used a DIP-seq approach to measure changes in 5hmC across the genome. 5hmC is a stable epigenetic modification and may influence transcription, and while the nature of this influence is unclear studies suggest that 5hmC enrichment at transcriptional start sites (TSSs) leads to repression of transcription (Wu et al., 2011). Recent studies in mice showed that 5hmC enrichment is reversibly altered at specific loci in response to fat accumulation in the liver, indicating that the TET enzymes may play a role in NAFLD pathogenesis (Lyall et al., 2020). We did not observe wide-scale changes in 5hmC enrichment, and 5hmC changes at specific loci did not, in general, correlate with transcriptional changes in steatotic cells, except at a limited number of promoter regions. As NAFLD comprises a spectrum of pathologies, it is possible that our model reflects a different disease stage or that the changes in other studies are a consequence of the use of whole tissue. It is also possible that differential 5hmC enrichment is occurring in regions associated with expression of non-coding RNAs, including long noncoding RNAs or microRNAs, which in turn may influence mRNA transcription (Hu et al., 2017; Wang et al., 2019a). Other mechanisms by which fumarate, as an allosteric inhibitor of the dioxygenase family of enzymes may impact on transcriptional regulation in NAFLD, include through effects on HIF1α or the histone demethylase enzymes (Han et al., 2019; Kim et al., 2018).

Taken together, we demonstrate for the first time that the development of steatosis in HLCs associates with truncation of the TCA cycle. We further demonstrate that on exposure to LPO, mitochondrial metabolic pathways are re-wired in order to maintain TCA cycle activity, through PNC-driven fumarate accumulation, which then results in the development of macrovesicular steatosis. These findings reveal a previously unknown mechanism linked to hepatic steatosis and may lead to further understanding of transcriptional and metabolic rewiring associated with NAFLD.

## Acknowledgements

MCS was supported by a British Heart Foundation PhD studentship (FS/16/54/32730) and by the British Heart Foundation Centre of Research Excellence. SW-Z was funded by the Deutsche Forschungsgemeinschaft (SFB960). This work was supported in part by the Wellcome Trust [grant number 208400/Z/17/Z] and we thank HWB-NMR at the University of Birmingham for providing open access to their Wellcome Trust-funded NMR equipment. AJD was funded by the British Heart Foundation Centre of Research Excellence, University of Edinburgh. Our thanks go to the Wellcome Trust Clinical Research Facility Genetics Core, Western General Hospital, Edinburgh, UK. DCH, JMR, BLV were supported with awards from the MRC Doctoral Training Partnership (MR/K501293/1) and the Chief Scientist Office (TCS/16/37).

## Data Accessibility

Transcriptomic sequencing data have been deposited on the GEO repository (accession number GSE138052). 5hmC sequencing data have also been deposited on the GEO repository (accession number GSE144955).

## Author Contributions

MCS was involved in conceptualisation, methodology, validation, formal analysis, investigation, data curation, writing (original draft preparation, reviewing and editing) and visualisation. BLV, JMR, PDW and AT were involved in methodology, validation, formal analysis, investigation and writing (reviewing and editing). SW-Z and JPT were involved in software, formal analyses, data curation and writing (reviewing and editing). CL and DAT were involved in conceptualisation, methodology, validation, formal analysis, investigation, data curation, provision of resources, writing (reviewing and editing), project administration, funding acquisition and supervision. DCH and AJD were involved in conceptualisation, methodology, validation, formal analysis, provision of resources, writing (reviewing and editing), project administration, funding acquisition and supervision.

## Materials and Methods

### Differentiation of pluripotent human stem cells to hepatocyte-like cells and induction of intracellular lipid accumulation

Human female H9 pluripotent stem cells (PSCs) were differentiated to hepatocyte-like cells (HLCs) as previously described (Wang et al., 2017). HLCs were cultured in a 96-well format for measurements of lipid accumulation and in a 6-well format for all other analyses. To assess loss of pluripotency and development of a gain of a hepatocyte-like phenotype, we measured expression of pluripotency (NANOG) and hepatocyte (ALB, HNF4A) markers throughout the differentiation process (Fig. S1A). Details of primers and Universal Probe Library probes (Roche) can be found in Table S1. CYP3A4 activity was similar to that previously observed, demonstrating that HLCs are functionally similar to hepatocytes (Wang et al., 2017), and this was not diminished by intracellular lipid accumulation (Fig. S1B). Lipid accumulation was induced as previously described (Lyall et al., 2018). Briefly, at day 17, HLCs were exposed to a cocktail of sodium l-lactate (L; 10mM), sodium pyruvate (P; 1 mM) and octanoic acid (O; 2 mM) (Sigma, Gillingham, UK) for a period of 48 h. For isotopic tracing studies, lactate was replaced with ^13^C_3_-lactate (CK Isotopes, CLM-1579-05). For mechanistic studies, HLCs were exposed to either 5-Aminoimidazole-4-carboxamide-1-β-D-ribofuranosyl 5′-monophosphate (AICAR; 1mM; Sigma-Aldrich, A1393-50MG), O-(Carboxymethyl)hydroxylamine hemihydrochloride (AOA; 100 μM ; Sigma-Aldrich, C13408-1G) or AICAR combined with monomethyl fumarate (50 μM; Sigma-Aldrich, 651419-1G) for the same duration as LPO.

### RNA-seq analysis

Total RNA was extracted from HLCs using the Monarch^®^ Total RNA Miniprep Kit (New England BioLabs, T2010). RNA integrity was assessed using a Bioanalyzer (Agilent) with the RNA 6000 Nano kit. All samples had a RIN value >7.0. mRNA sequencing was performed on 3 biological replicates per group by the Beijing Genomics Institute (BGI) (Shenzhen, China). Library preparation was performed with the TruSeq Stranded mRNA Library Preparation kit (Illumina, RS-122-2101), with additional use of the Ribo-Zero Gold rRNA Removal Kit (Illumina, MRZG12324). Paired-end sequencing was performed on an Illumina HiSeq 4000, with each sample sequenced to a depth >90 million reads. The generated FASTQ files were trimmed to remove adapters, using Trimmomatic (version 0.36) (Bolger et al., 2014), before performing quality control with FastQC (version 0.11.4) (Andrews). Alignment was performed against the *Homo sapiens* GRCh19 assembly. The assembly was first indexed using STAR (version 2.5.1b) before mapping trimmed reads, using STAR (version 2.5.1b) in paired-end mode with default behaviour (Dobin and Gingeras, 2015). Duplicate reads were removed using Picard (version 2.7.11) (2018), before using featureCounts to generate raw read counts for each gene. Differential gene expression (DEG) analysis was performed using DESeq2 (Love et al., 2014). Heatmaps were generated with Heatmapper (Babicki et al., 2016). Pathway enrichment analysis was performed using the Kyoto Encyclopedia of Genes and Genomes (KEGG) function (Kanehisa, 2019; Kanehisa and Goto, 2000; Kanehisa et al., 2019) of the Database for Annotation, Visualization, and Integrated Discovery (DAVID) (Huang et al., 2009a, 2009b).

### Real-time quantitative PCR

RNA was taken from that prepared for RNA-sequencing. cDNA was generated using the High Capacity cDNA Reverse Transcriptase Kit (Applied Biosystems, 4368814). A master mix was prepared using PerfeCTa FastMix II (Quanta Biosciences, Inc., 95118-250). cDNA was amplified and quantified using the Universal Probe Library (Roche, Burgess Hill, UK) system on a Roche LightCycler 480 (Roche Diagnostics Ltd, Switzerland).

### NMR Spectroscopy

This protocol was previously described by Hollinshead *et al* (Hollinshead et al., 2018). At the conclusion of tracer experiments, cells were washed with 2 mL ice-cold 0.9% saline solution and quenched with 0.3 mL pre-chilled methanol (−20 °C). After adding an equal volume of ice-cold HPLC-grade water containing 1 μg/mL D6-glutaric acid (C/D/N Isotopes Inc), cells were collected with a cell scraper and transferred to tubes containing 0.3 mL of chloroform (−20 °C). The extracts were shaken at 1400 rpm for 20 min at 4 °C and centrifuged at 16,000 × g for 5 min at 4 °C. Then, 0.3 mL of the upper aqueous phase was collected and evaporated in eppendorfs, under a vacuum using a Savant™ SpeedVac™ Concentrator (ThermoFisher). These samples were used either for NMR spectroscopy of for GC-MS. For NMR, dried samples were re-suspended in 60 μL of 100 mM sodium phosphate buffer (pH 7.0) containing 500 μM DSS and 2 mM Imidazole, 10% D20, pH 7.0. Samples were vortexed, sonicated (5-15 min) and centrifuged briefly, before transferred to 1.7 mm NMR tubes using an automated Gilson. One-dimensional (1D)-^1^H NMR spectra and two-dimensional (2D)-^1^H,^13^C Heteronuclear Single Quantum Coherence Spectroscopy (HSQC) NMR spectra were acquired using a 600 MHz Bruker Avance III spectrometer (Bruker Biospin) with an inverse cryogenic probe for 1.7 mm NMR sample tubes, fitted with a z-axis pulsed field gradient, at 300 K. Spectral widths were set to 13 and 160 ppm for the ^1^H and ^13^C dimensions, respectively. For the indirect (^13^C) dimension of the 2D-^1^H,^13^C HSQC NMR spectra, 1228 out of 4096 (30%) data points were acquired using a non-uniform sampling scheme. ^13^C-^13^C splittings were enhanced 4-fold in the ^13^C dimension. Each sample was automatically tuned, matched and then shimmed (1D-TopShim) to a DSS line width of <1 Hz before acquisition of the first spectrum. Total experiment time was ~15 min per sample for 1D-^1^H NMR spectra and 1 h per sample for 2D-^1^H,^13^C HSQC NMR spectra. 1D-^1^H NMR spectra were processed using the MATLAB-based MetaboLab software (Ludwig and Günther, 2011). All 1D data sets were apodized using a 0.3 Hz exponential window function and zero-filled to 131,072 data points before Fourier Transformation. The chemical shift was calibrated by referencing the DSS signal to 0 ppm. 1D-^1^H NMR spectra were manually phase corrected. Baseline correction was achieved using a spline function (Ludwig and Günther, 2011). 1D-^1^H-NMR spectra were exported into Bruker format for metabolite identification and concentration determination using Chenomx 7.0 (Chenomx INC). 2D-^1^H,^13^C HSQC NMR spectra were reconstructed using compressed sensing in the MDDNMR and NMRpipe software (Delaglio et al., 1995; Kazimierczuk and Orekhov, 2011; Orekhov and Jaravine, 2011). The final spectrum size was 922 real data points for the ^1^H dimension and 16,384 real data points for the ^13^C dimension. Analysis was performed using MetaboLab and pyGamma software was used in multiplet simulations (Smith et al., 1994). The methyl group of lactate was used to calibrate the chemical shift based on its assignment in the human metabolome database (Wishart et al., 2013).

### GC-MS

Dried polar metabolites were purified as described for NMR spectroscopy. These were derivatised by incubating with 40 μL 2% methoxyamine hydrochloride (Sigma Aldrich, 226904) in pyridine (Thermo Fisher Scientific, 25104) at 60 °C for 1 h, followed by incubation with 60 μL *N*-methyl-*N*-*tert*-butyldimethylsilyltrifluoroacetamide with 1% *tert*-butyldimethylchlorosilane (MTBSTFA with 1% t-BDMCS) at 60 °C for 1 h.

GC-MS analysis was performed using an Agilent 6890GC in combination with an Agilent 5975C MS. The MS was operated under electron impact ionization at 70 eV with the source held at 230 °C and the quadrupole at 150 °C. Helium was used as the carrier gas and maintained at a flow rate of 1 mL/min. 1 μL of derivatised sample was injected (splitless) with an inlet temperature of 280 °C on to a Rxi-5MS column (Restek) The oven temperature was held at 100 °C for 1 min then increased at a rate of 5 °C/min up to a maximum temperature of 330 °C. Ions were detected using selected ion monitoring (SIM) mode as previously described (Battello et al., 2016). MetaboliteDetector software was used to correct for the natural isotope distribution and to determine the mass isotopomer distribution (MID) (Hiller et al., 2009).

### DNA hydroxymethylation immunoprecipitation and sequencing (hmeDIP-sequencing)

DNA was purified using the Monarch^®^ Genomic DNA Purification kit (New England BioLabs, T3010S). DNA immunoprecipitation and sequencing was performed as previously described, using the Ion Proton platform (Thomson et al., 2015), with the addition of an IgG control (Merck, 12-370). We sequenced three biological replicates per group. A mean read length of 137-147 base pairs and 21,130,039 - 31,693,844 reads per sample was achieved. Reads were aligned to the hg19 genome using Torrent Suite v5.2.0. Aligned reads were sorted using SAMtools, before calling peaks using MACS2 (v. 2.1.1) -f BAM --broad --broad-cutoff 0.05 -B -g hs, over corresponding inputs (Zhang et al., 2008). To detect differentially hydroxymethylated regions (DHRs), we used Diffbind with DESeq2 (Stark and Brown). For Diffbind analysis, data were normalised to a pooled input for each group and an IgG control. DHMRs were assigned to genes and other genomic features using the HOMER (v. 4.8; hg19) annotatePeaks tools (Heinz et al., 2010). For candidate hmeDIP analysis, the concentration of each sample was extrapolated from a standard curve of arbitrary concentrations and normalised to 10% input. Regions of interest were identified from the hmeDIP-sequencing dataset. Primers were designed using the NCBI primer-BLAST software (Table S2). Data are available through the Gene Expression Omnibus (GSE144955). Sliding window profiles and heatmaps were generated using deepTools (Ramírez et al., 2014), using the plotProfile and plotHeatmap functions, respectively, with blacklisted regions subtracted.

### High content analysis microscopy

Cells were stained with a cell painter assay, adapted from Lyall *et al* and Bray *et al* (Bray et al., 2016; Lyall et al., 2018). Cells were fixed with 50 μL/well 4% (wt/vol) paraformaldehyde (Electron Microscopy Sciences, 15710-S) for 15 minutes at room temperature. For permeabilisation, cells were incubated in 0.1% Triton X-100 (Sigma-Aldrich, T8787) in PBS for 15 minutes at room temperature. For lipid droplet analysis, cells were then stained with a combination of NucBlue Live ReadyProbes^®^ Reagent (2 drops/mL) (Molecular Probes, R37605), HCS CellMask™ Red (2 μL/10 mL) (Invitrogen, H32712), and BODIPY™ 493/503 (1:1000) (Life Sciences, D3922), as per the manufacturer’s instructions. Following staining, images were acquired using an Operetta High Content Analysis microscope (Perkin Elmer, Buckinghamshire, UK). Lipid droplet morphology was analysed as previously described (Lyall et al., 2018).

### Statistical analysis

All statistical analyses were performed using Graph Prism Version 8.0 for Windows or macOS, GraphPad Software, La Jolla California USA, www.graphpad.com. Normality of data distribution was measured using the Shapiro-Wilks test. Where indicated, data were analysed by unpaired Student’s t-test, Mann-Whitney test, one-way analysis of variance (ANOVA) or two-way ANOVA. Data were considered to be significant where *p* <0.05.

