## Supplemental files and tables for "Macrovesicular steatosis in nonalcoholic fatty liver disease is a consequence of purine nucleotide cycle driven fumarate accumulation" (from another version of this preprint: version 1, 2020-06-24)

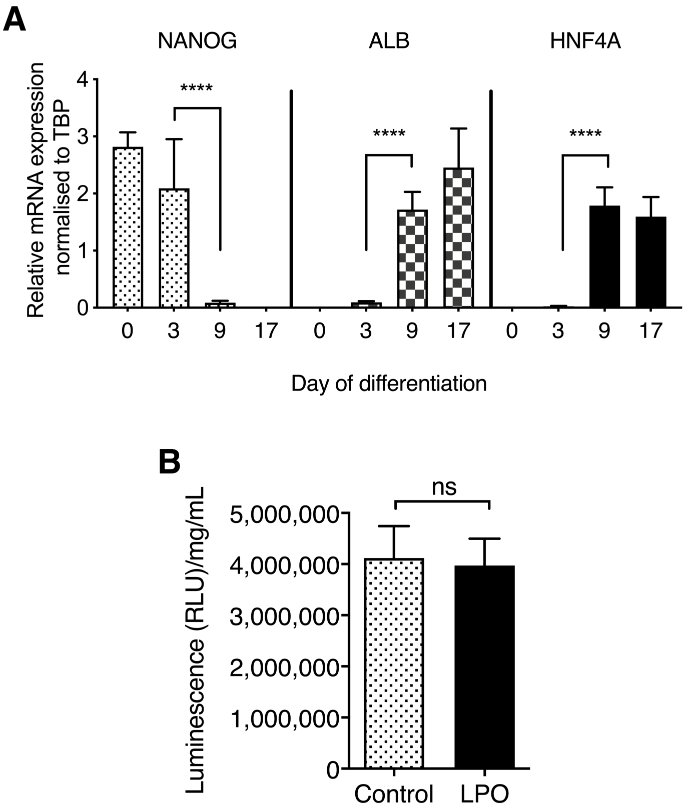


**Figure S1.** Validation of H9 to HLC differentiation process. (**A**) Between days 0 and 17 of differentiation, H9 cells lose expression of the pluripotency transcript NANOG, but gain expression of two hepatocyte transcripts, ALB and HNF4A. (**B**) HLCs are functionally active, demonstrating robust CYP4A4 activity, which is not diminished following induction of steatosis. Expression groups had *n* = 3 biological replicates/group. CYP3A4 *n* = 8 biological replicates/group. Data were analysed by analysed by (**A**) two-way ANOVA with Sidak post-hoc testing or (**B**) two-tailed Student's t-test. Data are expressed as mean ± SD.

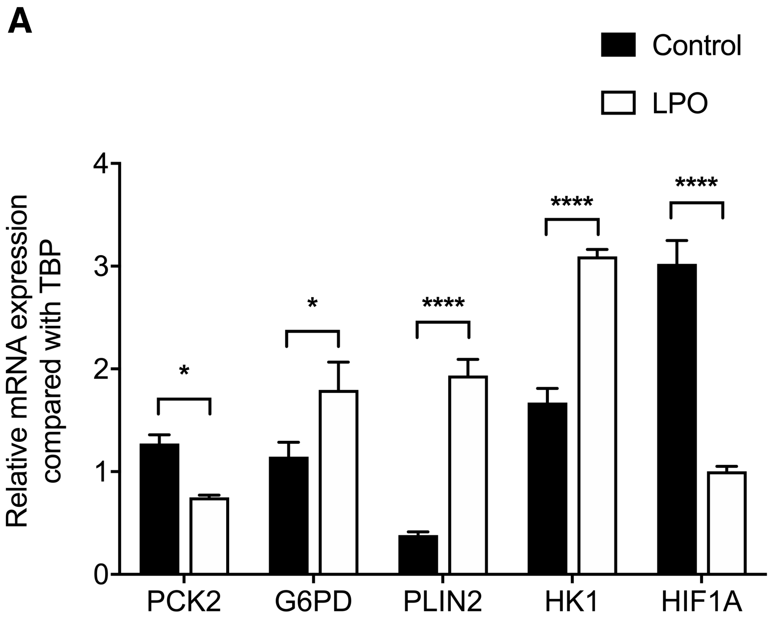


**Figure S2.** Validation of mRNA sequencing analysis by RT-qPCR. (**A**) RT-qPCR analysis of key genes validated mRNA sequencing data. Data were analysed by analysed by two-way ANOVA with Sidak post-hoc testing. Data are expressed as mean ± SD.

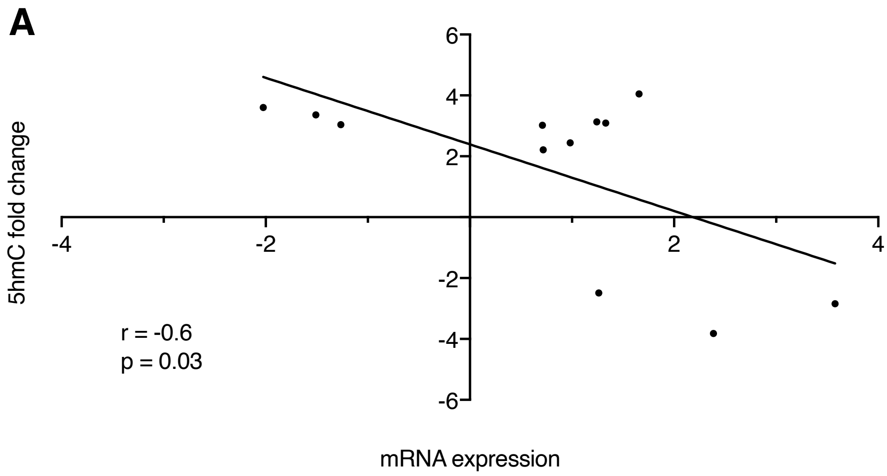


**Figure S3.** Steatotic HLCs show a moderate negative correlation between promoter 5hmC enrichment and mRNA expression. The cut-off for inclusion was: 5hmC) Enrichment/depletion >2.0; mRNA expression) $Log_2$ fold change >0.5. Data were analysed by linear regression.

**Table S1. Primers sequences for RT-qPCR**

| Gene | Forward sequence | Reverse sequence | Probe |
|---|---|---|---|
| TBP | GAACATCATGGATCAGAACAACA | ATAGGGATTCCGGGAGTCAT | 87 |
| NANOG | ATGCCTCACACGGAGACTGT | CAGGGCTGTCCTGAATAAGC | 69 |
| ALB | GTGAGGTTGCTCATCGGTTT | GAGCAAAGGCAATCAACACC | 7 |
| HNF4A | AGCAACGGACAGATGTGTGA | TCAGACCCTGAGCCACCT | 27 |
| PCK2 | CGAAAGCTCCCCAAGTACAA | GCTCTCTACTCGTGCCACATC | 20 |
| G6PD | AACAGAGTGAGCCCTTCTTCA | GGAGGCTGCATCATCGTACT | 5 |
| PLIN2 | TCAGCTCCATTCTACTGTTCACC | CCTGAATTTTCTGATTGGCACT | 72 |
| HK1 | GACCAAGTTTCTCTCTCAGATCG | CCTAGCTGCTGGAGGATAGC | 1 |
| HIF1A | GATAGCAAGACTTTCCTCAGTCG | TGGCTCATATCCCATCAATTC | 64 |

**Table S2. Primer sequences for validation of DNA immunoprecipitation protocol**

| Gene | Forward sequence | Reverse sequence |
|---|---|---|
| GAPDH promoter – negative control | CGGCTACTAGCGGTTTTACG | AAGAAGATGCGGCTGACTGT |
| H19 genic – positive control | GATCTCGGCCCTAGTGTGAA | GTGATGTGTGAGCCTGCACT |
| UBIAD1 genic – positive control | CTCTTCCTCCTCCTCGTCCT | CATCCAGGAACCACAGTCCT |

**Table S3. Upregulated and downregulated KEGG pathways**

| **Pathways with upregulated genes** | | |
|---|---|---|
| *Pathway* | *Benjamini* | *Genes* |
| hsa04713: Circadian entrainment | 4.20 | ADCY4, ADCY2, CACNA1I, GRIN1, GRIA3, PRKG1, PRKCB, KCNJ5, PLCB4, GRIN2D, CACNA1G, RYR1, GNG2, PER3, GNG4, MTNR1A |
| hsa04724: Glutamatergic synapse | 4.15 | ADCY4, DLGAP1, ADCY2, GRIK2, GRIN1, GRIK5, GRIN3B, GRIA3, SHANK1, PRKCB, SLC17A7, GLS2, GRM4, PLCB4, GRIN2D, PLA2G4F, GNG2, GNG4, PLA2G4D |
| hsa04725: Cholinergic synapse | 3.59 | ADCY4, ACHE, ADCY2, KCNJ12, KCNJ14, PRKCB, KCNJ4, KCNQ4, KCNQ3, CHRM4, PLCB4, GNG2, PIK3R5, GNG4, KCNQ1, CHRNA3 |
| hsa04750: Inflammatory mediator regulation of TRP channels | 3.31 | ADCY4, ADCY2, TRPM8, TRPV2, ASIC3, ASIC1, PRKCB, PLCB4, PLA2G4F, PIK3R5, ALOX12, NGF, PLA2G4D |
| hsa05414: Dilated cardiomyopathy | 3.26 | ADCY4, ADCY2, ADRB1, SGCG, CACNG8, ITGA8, ITGA7, CACNB1, CACNB4, TNNI3, ITGA2B |
| hsa04911: Insulin secretion | 3.23 | TRPM4, ADCY4, ADCY2, PLCB4, KCNN1, KCNN3, ATP1A3, RIMS2, PCLO, KCNJ11, PRKCB |
| hsa04726: Serotonergic synapse | 3.14 | GABRB2, PRKCB, KCNJ5, PLCB4, ALOX15B, HTR7, SLC18A2, PLA2G4F, GNG2, HTR1D, HTR3A, GNG4, ALOX12, PLA2G4D |
| hsa04514: Cell adhesion molecules (CAMs) | 2.81 | ICAM1, NTNG1, NTNG2, CLDN10, CD40, HLA-DMB, CDH4, HLA-G, NCAM2, ITGA8, NLGN4X, PECAM1, CNTN1, HLA-DOA, CD6, ICOSLG |
| hsa04921: Oxytocin signalling pathway | 2.66 | ADCY4, ADCY2, CACNG8, CACNB1, CACNB4, KCNJ12, TRPM2, KCNJ14, PRKCB, KCNJ5, KCNJ4, PLCB4, RYR1, PLA2G4F, NFATC1, PLA2G4D |
| hsa04080: Neuroactive ligand-receptor interaction | 2.43 | F2RL2, C5AR1, GABRB2, GRIK2, GRIN1, GABBR1, GRIK5, LPAR3, GRIA3, GRIN3B, P2RX5, GRM4, SSTR2, ADRB1, S1PR1, CHRM4, CHRNA9, SSTR1, GRIN2D, HTR7, CHRNA5, ADRA1A, CALCRL, HTR1D, CHRNA3, GRID1, MTNR1A |
| hsa04024: cAMP signalling pathway | 2.39 | ADCY4, HCN2, ADCY2, GRIN1, GABBR1, ATP1A3, GRIN3B, GRIA3, TNNI3, GLI1, SSTR2, ADRB1, SSTR1, PDE4A, GRIN2D, PIK3R5, HTR1D, HCAR2, NFATC1 |

| | | |
|---|---|---|
| hsa04020: Calcium signalling pathway | 2.37 | SLC8A3, ADCY4, SLC8A2, ADCY2, CACNA1I, GRIN1, PRKCB, P2RX5, PLCB4, ADRB1, ATP2A3, GRIN2D, HTR7, CACNA1G, RYR1, ADRA1A, PLCD1 |
| hsa04014: Ras signalling pathway | 2.32 | FGF19, FGF5, FLT4, FGF17, EFNA3, GRIN1, FGF11, PRKCB, RASAL1, HTR7, RASGRP2, PLA2G4F, PLA1A, GNG2, PIK3R5, NGFR, SYNGAP1, GNG4, RASA4, PLA2G4D, NGF |
| **Pathways with downregulated genes** | | |
| *Pathway* | *Benjamini* | *Genes* |
| hsa05322: Systemic lupus erythematosus | 6.65 | HIST1H2AB, C7, HIST1H4L, HIST1H2AG, C6, HIST1H2AE, HIST1H2BO, HIST2H2AB, HIST1H2BM, HIST1H4A, HIST1H2BL, HIST1H2BI, HIST2H2AC, HIST1H2BJ, H2AFX, HIST3H2BB, HIST1H4I, HIST1H4J, HIST2H3A, HIST1H3J, HIST1H2BB, HIST1H2BC, HIST1H2BE, HIST1H2BF, HIST1H2BG, HIST1H2BH, ACTN2, HIST2H3C, HIST2H3D, HIST2H2BF, HIST1H3A, HIST1H3B, HIST1H2AI, HIST1H2AH, HIST1H3C, HIST1H3E, HIST1H2AJ, HIST1H3F, HIST1H2AM, HIST1H3G, HIST1H2AL, HIST1H3H, HIST1H3I |
| hsa00140: Steroid hormone biosynthesis | 6.43 | HSD3B2, CYP3A4, CYP3A5, CYP3A7, HSD3B1, CYP11A1, HSD17B1, UGT1A9, CYP7A1, UGT2B11, UGT2B4, HSD17B6, UGT2B10, UGT2A3, SULT1E1, UGT2B15, AKR1D1, CYP19A1 |
| hsa00053: Ascorbate and aldarate metabolism | 5.37 | UGT1A9, UGT2B11, RGN, UGT2B4, UGT2B10, UGT2A3, UGT2B15 |
| hsa04610: Complement and coagulation cascades | 5.10 | KNG1, F11, PLAT, MBL2, C7, MASP2, F13A1, C6, F9, C4BPB, C4BPA, F13B, F3, KLKB1, SERPINA5, SERPIND1, CPB2 |
| hsa00040: Pentose and glucuronate interconversions | 5.02 | UGT1A9, KL, AKR1B10, UGT2B11, UGT2B4, UGT2B10, UGT2A3, UGT2B15 |
| hsa05034: Alcoholism | 4.92 | HIST1H2AB, HIST1H4L, HIST1H2AG, HIST1H2AE, HIST1H2BO, HIST2H2AB, HIST1H2BM, HIST1H4A, HIST1H2BL, HIST1H2BI, HIST2H2AC, HIST1H2BJ, H2AFX, HIST3H2BB, HIST1H4I, HIST1H4J, HIST2H3A, HIST1H3J, HIST1H2BB, HIST1H2BC, HIST1H2BE, HIST1H2BF, HIST1H2BG, HIST1H2BH, FOSB, HIST2H3C, HIST2H3D, HIST2H2BF, NTRK2, HIST1H3A, |

| | | |
|---|---|---|
| | | HIST1H3B, HIST1H2AI, HIST1H2AH, HIST1H3C, HIST1H3E, HIST1H2AJ, HIST1H3F, HIST1H2AM, HIST1H3G, HIST1H2AL, HIST1H3H, HIST1H3I |
| hsa00982: Drug metabolism - cytochrome P450 | 4.88 | GSTA1, CYP3A4, GSTA2, CYP3A5, ALDH3B2, ADH1B, ADH1A, FMO5, UGT1A9, FMO1, ADH4, UGT2B11, UGT2B4, UGT2A3, UGT2B10, UGT2B15 |
| hsa00830: Retinol metabolism | 4.86 | CYP3A4, CYP3A5, UGT1A9, CYP3A7, ADH4, UGT2B11, ADH1B, UGT2B4, CYP26A1, HSD17B6, ADH1A, UGT2B10, UGT2A3, UGT2B15, RDH5 |
| hsa05204: Chemical carcinogenesis | 4.40 | GSTA1, CYP3A4, GSTA2, CYP3A5, CYP3A7, NAT2, ADH1B, ALDH3B2, ADH1A, CYP3A43, UGT1A9, ADH4, UGT2B11, UGT2B4, UGT2A3, UGT2B10, UGT2B15 |
| hsa00983: Drug metabolism - other enzymes | 4.05 | CYP3A4, UGT1A9, NAT2, UGT2B11, UGT2B4, UGT2B10, UGT2A3, UGT2B15, TK1 |
| hsa00980: Metabolism of xenobiotics by cytochrome P450 | 3.92 | GSTA1, CYP3A4, CYP3A5, GSTA2, ALDH3B2, ADH1B, ADH1A, UGT1A9, ADH4, UGT2B11, UGT2B4, UGT2A3, UGT2B10, UGT2B15 |
| hsa04110: Cell cycle | 3.01 | CDC6, CDK1, PKMYT1, TTK, CDC20, ESPL1, PTTG1, MCM2, CDC25C, CCNB1, CDKN1C, MAD2L1, CCNB2, PLK1, CDKN2C, BUB1, BUB1B, CCNA2 |
| hsa04114: Oocyte meiosis | 2.61 | CCNB1, CDK1, MAD2L1, CCNB2, PLK1, SGO1, BUB1, FBXO43, PKMYT1, AURKA, ESPL1, CDC20, PTTG1, CDC25C |
| hsa05202: Transcriptional misregulation in cancer | 2.36 | PLAT, HIST2H3A, NFKBIZ, HIST1H3J, MMP9, MMP3, HIST2H3C, HIST2H3D, HHEX, EYA1, CDKN2C, HIST1H3A, HIST1H3B, HIST1H3C, HIST1H3E, HIST1H3F, HIST1H3G, HIST1H3H, HIST1H3I |

**Table S4. Dysregulated genes in the TCA Cycle KEGG pathway**

| *Ensembl ID* | *Gene* | *$Log_2FC$* | *padj* |
|---|---|---|---|
| ENSG00000197444 | OGDHL | 1.08547769 | 3.29E-30 |
| ENSG00000166411 | IDH3A | 0.74709247 | 7.70E-20 |
| ENSG00000204370 | SDHD | 0.3777972 | 0.0003 |
| ENSG00000100412 | ACO2 | 0.26021698 | 0.002 |
| ENSG00000146701 | MDH2 | 0.23945724 | 0.0009 |
| ENSG00000073578 | SDHA | -0.1593416 | 0.03 |
| ENSG00000138413 | IDH1 | -0.1812103 | 0.03 |
| ENSG00000168291 | PDHB | -0.2026402 | 0.02 |
| ENSG00000131828 | PDHA1 | -0.2107741 | 0.01 |
| ENSG00000163541 | SUCLG1 | -0.2111892 | 0.006 |
| ENSG00000143252 | SDHC | -0.2414483 | 0.02 |
| ENSG00000150768 | DLAT | -0.2619025 | 0.0003 |
| ENSG00000091483 | FH | -0.4936542 | 1.01E-12 |
| ENSG00000014641 | MDH1 | -0.5085681 | 3.36E-13 |
| ENSG00000101365 | IDH3B | -0.5405112 | 5.07E-13 |
| ENSG00000067829 | IDH3G | -0.5829934 | 1.14E-09 |
| ENSG00000100889 | PCK2 | -0.5852916 | 3.07E-18 |
| ENSG00000131473 | ACLY | -0.6074454 | 9.80E-24 |
| ENSG00000173599 | PC | -0.9409933 | 3.13E-37 |
| ENSG00000182054 | IDH2 | -1.0677314 | 8.52E-45 |

**Table S5. Dysregulated genes in the Oxidative Phosphorylation KEGG pathway**

| *Ensembl ID* | *Gene* | *$Log_2FC$* | *padj* |
|---|---|---|---|
| ENSG00000147614 | ATP6V0D2 | 3.973933876 | 9.35E-140 |
| ENSG00000198763 | ND2 | 1.473526224 | 2.53E-06 |
| ENSG00000198888 | ND1 | 1.363322749 | 6.19E-05 |
| ENSG00000198840 | ND3 | 1.217622162 | 2.54E-33 |
| ENSG00000212907 | ND4L | 1.095495883 | 0.0001 |
| ENSG00000198886 | ND4 | 0.866690501 | 0.002 |
| ENSG00000114573 | ATP6V1A | 0.803262336 | 2.80E-32 |
| ENSG00000198804 | COX1 | 0.656245362 | 1.17E-09 |
| ENSG00000198899 | ATP6 | 0.634543196 | 0.009 |
| ENSG00000198938 | COX3 | 0.632275024 | 2.80E-10 |
| ENSG00000171130 | ATP6V0E2 | 0.589272395 | 2.52E-11 |
| ENSG00000198727 | CYTB | 0.557687429 | 0.02 |
| ENSG00000047249 | ATP6V1H | 0.506014583 | 5.94E-13 |
| ENSG00000117410 | ATP6V0B | 0.479535962 | 2.58E-11 |
| ENSG00000147416 | ATP6V1B2 | 0.437789019 | 5.51E-09 |
| ENSG00000128609 | NDUFA5 | 0.381648881 | 0.0008 |
| ENSG00000204370 | SDHD | 0.377797199 | 0.0003 |
| ENSG00000198712 | COX2 | 0.322901415 | 0.005 |
| ENSG00000073578 | SDHA | -0.159341647 | 0.04 |
| ENSG00000023228 | NDUFS1 | -0.210017339 | 0.002 |
| ENSG00000112695 | COX7A2 | -0.211631763 | 0.02 |
| ENSG00000156467 | UQCRB | -0.2162753 | 0.03 |
| ENSG00000167792 | NDUFV1 | -0.222597617 | 0.007 |
| ENSG00000135390 | ATP5MC2 | -0.222831627 | 0.002 |
| ENSG00000143252 | SDHC | -0.24144827 | 0.02 |
| ENSG00000160194 | NDUFV3 | -0.242343108 | 0.004 |
| ENSG00000131100 | ATP6V1E1 | -0.274149444 | 2.61E-05 |
| ENSG00000189043 | NDUFA4 | -0.277227615 | 0.0001 |
| ENSG00000176340 | COX8A | -0.278696825 | 0.001 |
| ENSG00000165264 | NDUFB6 | -0.28300413 | 0.005 |
| ENSG00000213619 | NDUFS3 | -0.29075047 | 0.001 |
| ENSG00000178741 | COX5A | -0.297956503 | 9.27E-05 |
| ENSG00000110719 | TCIRG1 | -0.300385483 | 0.002 |
| ENSG00000090266 | NDUFB2 | -0.301527793 | 0.001 |
| ENSG00000174886 | NDUFA11 | -0.318594422 | 0.001 |
| ENSG00000169021 | UQCRFS1 | -0.319382279 | 2.33E-05 |
| ENSG00000130414 | NDUFA10 | -0.320916351 | 3.89E-05 |
| ENSG00000131143 | COX4I1 | -0.336149045 | 2.75E-07 |
| ENSG00000158864 | NDUFS2 | -0.343567449 | 4.23E-06 |
| ENSG00000139180 | NDUFA9 | -0.357605693 | 0.0006 |
| ENSG00000004779 | NDUFAB1 | -0.360587197 | 2.52E-05 |
| ENSG00000184076 | UQCR10 | -0.375287362 | 3.00E-06 |
| ENSG00000116459 | ATP5PB | -0.379803823 | 9.50E-08 |
| ENSG00000126267 | COX6B1 | -0.388197692 | 3.84E-08 |
| ENSG00000100554 | ATP6V1D | -0.405728961 | 7.75E-08 |
| ENSG00000131495 | NDUFA2 | -0.419069805 | 1.65E-06 |

| | | | |
|---|---|---|---|
| ENSG00000115286 | NDUFS7 | -0.422028053 | 0.0002 |
| ENSG00000136888 | ATP6V1G1 | -0.428892928 | 2.02E-09 |
| ENSG00000183648 | NDUFB1 | -0.432392964 | 0.0003 |
| ENSG00000154518 | ATP5MC3 | -0.445435152 | 1.00E-08 |
| ENSG00000125356 | NDUFA1 | -0.459056555 | 2.84E-07 |
| ENSG00000179091 | CYC1 | -0.468189203 | 1.54E-09 |
| ENSG00000127540 | UQCR11 | -0.475454692 | 4.04E-09 |
| ENSG00000110955 | ATP5F1B | -0.49393262 | 1.83E-16 |
| ENSG00000099795 | NDUFB7 | -0.509075592 | 1.10E-09 |
| ENSG00000168653 | NDUFS5 | -0.520875625 | 8.23E-11 |
| ENSG00000159720 | ATP6V0D1 | -0.540902137 | 6.67E-12 |
| ENSG00000147123 | NDUFB11 | -0.542107011 | 1.51E-08 |
| ENSG00000178127 | NDUFV2 | -0.542969651 | 0.006 |
| ENSG00000124172 | ATP5F1E | -0.574462256 | 1.96E-13 |
| ENSG00000152234 | ATP5F1A | -0.606022757 | 1.10E-17 |
| ENSG00000140990 | NDUFB10 | -0.613088113 | 6.15E-12 |
| ENSG00000119013 | NDUFB3 | -0.65837698 | 2.66E-11 |
| ENSG00000169020 | ATP5ME | -0.663226048 | 5.18E-18 |
| ENSG00000159199 | ATP5MC1 | -0.712677613 | 1.05E-19 |
| ENSG00000241468 | ATP5MF | -0.735984185 | 1.21E-15 |
| ENSG00000099624 | ATP5F1D | -0.737422752 | 3.56E-17 |
| ENSG00000169429 | CXCL8 | -0.820462236 | 3.09E-17 |
| ENSG00000164405 | UQCRQ | -0.860634282 | 1.66E-38 |
| ENSG00000198695 | ND6 | -0.979723775 | 1.42E-08 |
| ENSG00000143882 | ATP6V1C2 | -1.543440061 | 0.005 |

**Table S6. Promoter regions of genes with both altered mRNA expression and enrichment of 5hmC**

| Ensembl ID | Gene symbol | mRNA expression fold change | 5hmC fold change |
|---|---|---|---|
| ENSG00000181418 | DDN | 3.57 | -2.84 |
| ENSG00000105131 | EPHX3 | 2.38 | -3.82 |
| ENSG00000086619 | ERO1B | 1.65 | 4.05 |
| ENSG00000147408 | CSGALNACT1 | 1.33 | 3.09 |
| ENSG00000149927 | DOC2A | 1.26 | -2.49 |
| ENSG00000142156 | COL6A1 | 1.24 | 3.13 |
| ENSG00000137752 | CASP1 | 0.98 | 2.44 |
| ENSG00000119900 | OGFRL1 | 0.71 | 2.21 |
| ENSG00000145247 | OCIAD2 | 0.71 | 3.02 |
| ENSG00000183044 | ABAT | -1.26 | 3.04 |
| ENSG00000144395 | CCDC150 | -1.50 | 3.36 |
| ENSG00000167874 | TMEM88 | -2.02 | 3.60 |
